# Isolation and characterization of a highly specific monoclonal antibody targeting the botulinum neurotoxin type E exposed SNAP-25 neoepitope

**DOI:** 10.1101/2021.09.16.460610

**Authors:** Adva Mechaly, Eran Diamant, Ron Alcalay, Alon Ben-David, Eyal Dor, Amram Torgeman, Ada Barnea, Meni Girshengorn, Lilach Levin, Eyal Epstein, Ariel Tennenhouse, Sarel J. Fleishman, Ran Zichel, Ohad Mazor

## Abstract

Botulinum neurotoxin type E (BoNT/E), the fastest acting toxin of all BoNTs, cleaves the 25 kDa synaptosomal associated protein (SNAP-25) in motor neurons, leading to flaccid paralysis. Specific detection and quantification of BoNT/E-cleaved SNAP-25 neoepitope is essential for diagnosis of BoNT/E intoxication as well as for characterization of anti-BoNT/E antibody preparations. In order to isolate highly specific monoclonal antibodies suitable for in vitro immuno-detection of the exposed neoepitope, mice and rabbits were immunized with an eight amino acid peptide composed of the C-terminus of the cleaved SNAP-25. Immunized rabbits developed a specific and robust polyclonal antibody response, whereas immunized mice mostly demonstrated a weak antibody response that could not discriminate between the two forms of SNAP-25. An immune scFv phage-display library was constructed from the immunized rabbits and a panel of antibodies was isolated. Sequence alignment of the isolated clones revealed high similarity between both heavy and light chains, with exceptionally short HCDR3 sequences. A chimeric scFv-Fc antibody was further expressed and characterized, exhibiting a selective, ultra-high affinity (pM) towards the SNAP-25 neoepitope. Moreover, this antibody enabled sensitive detection of the cleaved SNAP-25 in BoNT/E treated SiMa cells with no cross reactivity with the intact SNAP-25. This novel antibody can be further used to develop an in vitro cell-based assay to diagnose BoNT/E intoxication and to characterize antitoxin preparations, thus eliminating the use of animals in the standard mouse bioassay.

## Introduction

Botulinum neurotoxins (BoNTs), the most potent toxins known in nature, are synthesized by the anaerobic bacterium *Clostridium botulinum* [1]. BoNT serotypes A, B, E, and rarely F are the causative agents of human botulism, a life-threatening disease [2]. BoNT is a single ∼150 kDa polypeptide, comprised of a ∼100 kDa heavy chain (HC) bridged by an S-S bond to a ∼50 kDa light chain (LC) which forms a zinc-dependent endopeptidase [3]. The C-terminal portion of the HC (H_C_) binds specific receptors on the presynaptic nerve ending membrane of cholinergic neurons, and the N-terminus of the heavy chain (H_N_) facilitates the translocation of the LC to the cytosol where it cleaves one of three soluble N-ethylmaleimide-sensitive factor attachment protein receptor (SNARE) proteins [4]. BoNT serotypes bind different receptors, and each serotype has a specific SNARE cleavage site, preventing the release of the neurotransmitter acetylcholine from nerve cells into the synapses [5-7]. Serotypes A and E cleave the 25 kDa synaptosomal associated protein (SNAP-25), serotypes B, D, F, and G cleave vesicle associated membrane protein (VAMP or synaptobrevin), and serotype C acts on both SNAP-25 and syntaxin [8]. Specifically, BoNT/E cleaves the 206 amino acid SNAP-25 protein (SNAP-25_1-206_) between Arg180 and Ile181 and as a result a truncated SNAP-25_1-180_ is formed [9, 10]. BoNT/E exerts its toxicity faster and significantly earlier than BoNT/A and BoNT/B requiring earlier diagnostic and medical responses.

Standard therapy for botulism in adults, including in cases of BoNT/E intoxications, involves treatment with antitoxin preparations [12, 13] derived from hyperimmune horses (A human antitoxin preparation is also available for infant botulism cases involving BoNT/A or BoNT/B [14]). The potencies of both the pharmaceutical antitoxin and the toxicity of BoNT/E preparations intended for horse immunization, are determined in vivo by the pharmacopeia mouse neutralization assay (PMNA) [15] and by the mouse bioassay (MBA) [16], respectively. However, these in vivo assays require a large number of laboratory animals, and hence development of alternative in vitro methods to measure activity of BoNT/E and concentration of neutralizing antibodies is imperative.

Assays with potential to replace MBA and PMNA should be based on in vitro demonstration of the three intoxication steps: receptor binding by the Hc, internalization by the HN, and enzymatic activity of the LC. Several methods can be applied to simulate the LC enzymatic activity in vitro using a synthetic substrate, and to further detect the resultant cleavage products by using either mass spectrometry, fluorescence, or an immunoassay with specific antibodies [17-23]. Yet, cell-based in vitro assays can combine the enzymatic activity step with the two earlier intoxication steps [24, 25]. Such assays are based on the correlation between the BoNT concentration incubated with the target cells and the specific quantitation of toxin cleavage product, detected using a specific antibody. Indeed, we have previously reported on a SiMa cell-based neutralization assay to measure the potency of anti-BoNT/A pharmaceutical antibody preparations in vitro [25]. This assay is based on a monoclonal chimeric antibody (MAb) that specifically recognizes the BoNT/A-cleaved SNAP-25 (anti SNAP-25_1-197_). Similarly, a proof-of-concept for the development of a SiMa cell-based neutralization assay to measure the activity of BoNT/E-cleaved SNAP-25 was shown before, using polyclonal antibodies [26]. Yet, in order to develop a reproducible assay, it is advisable to use well characterized monoclonal antibodies.

So far and despite the importance of developing such antibodies, only two groups have succeeded in isolating such antibodies using the traditional hybridoma cloning technique [27, 28]. We have previously isolated a panel of potent antibodies from immunized animals, by incorporating immunization methodologies that promote high affinity antibodies in vivo, together with efficient screening methods using phage-display libraries [29-31]. We thus hypothesized that, by applying similar methodologies, we would be able to isolate high-affinity antibodies that specifically recognize the cleaved SNAP-25 (SNAP-25_1-180)_ and not the intact SNAP-25_1-206_. Such antibodies could serve as the basis for sensitive in vitro neutralization assays. Here, we report the immunization strategy and selection procedures taken to reach this end and characterize the novel monoclonal antibody isolated.

## Materials and methods

### Peptides

All the peptides used in this work were synthesized by GenScript Japan Inc. (Tokyo, Japan) and are listed in table 1.

**Table 1:** Peptide sequences.

| Peptide | Sequence | Steps |
| --- | --- | --- |
| KLH-P1 | KLH-CTQNRQIDR | Rabbit immunization |
| Bio-P1 | Biotin-TQNRQIDR | 1. Monitoring serum titers<br>2. Library enrichment |
| Bio-P2 | Biotin-MGNEIDTQNRQIDRIMEKAD | Specificity (of serum and monoclonal Ab) |
| P3 | IIGNLRHMALDMGNEIDTQNRQIDRIMEKAD | Negative selection during library enrichment |

### Animal Immunization

Experiments were approved by the IIBR Animal Care and Use Committee and were conducted in accordance with the guidelines of the care and use of laboratory animals published by the Israeli Ministry of Health (protocols RB-27-16 and M-60-16). All efforts were made to minimize animal suffering. All animals were observed for morbidity and mortality, overt signs of toxicity, and any signs of distress throughout the study. Two female New Zealand White (NZW) rabbits (Oryctolagus cuniculus) and five female BALB/c mice (Mus musculus) were immunized with KLH-P1, a synthetic peptide corresponding to residues 173-180 of the human SNAP-25 sequence, conjugated to keyhole limpet hemocyanin (KLH). A cysteine residue, added to the N-terminus of the peptide, allowed coupling to KLH for immunization. Rabbits and mice were injected subcutaneously (s.c.) with 400 µg and 5 µg of antigen per animal, respectively, mixed either with Complete Freund’s Adjuvant (CFA) for priming or Incomplete Freund’s Adjuvant (IFA) for four booster immunizations.

### ScFv Library Construction and Screening

RNA from both rabbits was extracted from lymph nodes and spleen using the RNeasy mini kit and from blood samples using the RNeasy Protect Animal Blood kit (Qiagen GmbH, Hilden, Germany) according to the manufacturer′s instructions. All RNA samples were mixed together (per rabbit), and first-strand synthesis was performed using the Verso cDNA synthesis kit (Thermo Scientific, Waltham, MA, USA), with random hexamers and 1 µg RNA. VH and V_κ_ fragment amplification, construction of the scFv library and panning were carried out as described in extensive detail [29] with the following changes: A Maxisorp 96-well microtiter plate (Nunc, Sigma-Aldrich, St. Louis, MO, USA) was coated with 5 µg/ml streptavidin (Sigma s0977) diluted in NaHCO_3_ buffer (50 mM, pH 9.6) and incubated overnight. The plate was then washed, blocked (3% BSA + 0.05% tween20 in PBS) and loaded with synthetic peptide Bio-P1 (2 µg/ml). After a 30-minute incubation and a wash step, approximately 1 × 10^11^ pfu of blocked (3% skim milk + 0.05% tween20 in PBS) phage clones were incubated for 60 minutes with the peptide coated plate. In one of the enrichment schemes, the phage blocking buffer contained synthetic peptide P3 (5µg/ml). The plate was then washed once with blocker solution followed by a total of six washes with PBST (PBS, 0.05% Tween 20). Bound phage clones were then eluted and used to infect 5 mL of logarithmic-phase TG1 strain *E. coli* (Lucigen, Middleton, WI, USA). Two additional panning rounds were conducted (for the 2nd and 3rd rounds respectively) using 10^10^ and 10^9^ phage clones as input, while antigen incubation time was reduced to 30 and 15 min, phage blocking buffer was alternated (between 3% BSA to 3% Skim milk in PBS) and the PBST washing steps were raised to include 9 or 15 washes with PBST (0.1%).

### Production of Chimeric Antibodies

Phagemid DNA of the desired clones was isolated using QIAprep spin Miniprep kit (Qiagen, GmbH, Hilden, Germany), and the entire scFv was cloned into a mammalian immunoglobulin-based expression vector [32]. The vector was modified based on a published work [33], providing the scFv with the human constant Hc gene (IgG1), resulting in chimeric rabbit-human Fc-scFv antibody. FreeStyle Max 293 cells (Thermo Scientific, Waltham, MA, USA) were transiently transfected with the vector, and after a week, the supernatant was collected and the antibodies were purified on a HiTrap Protein-A column (GE healthcare, Little Chalfont, UK).

### ELISA

Maxisorp 96-well microtiter plates were coated overnight with 5 µg/mL streptavidin (50 µL/well) in NaHCO_3_ buffer (50 mM, pH 9.6), washed and blocked with PBST buffer (0.05% Tween 20, 2% BSA in PBS) for one hour. Synthetic primers Bio-P1 or Bio-P2 (2 µg/ml) were then loaded on the plate for 30 minutes. Rabbit sera, individual phage clones or purified antibodies were added to the plates for a one-hour incubation; the plates were then washed with PBST and incubated with the detecting antibody: alkaline phosphatase (AP)-conjugated goat anti rabbit for rabbit serum (Sigma-A8025), horseradish peroxidase (HRP)-conjugated anti-M13 antibody (GE healthcare, Little Chalfont, UK) for phage clones and AP-conjugated anti-human IgG (Sigma-A3187) for Fc-scFv antibodies. Detection of HRP conjugates was carried out with 3,3′,5,5′-tetramethybenzidine (TMB/E, Millipore, Billerica, MA, USA). Detection of AP conjugated antibodies was carried out with SIGMAFAST p-nitrophenyl phosphate tablets (Sigma-N2770).

### Affinity and Specificity Measurements

Binding studies were carried out using the Octet Red system (ForteBio, Version 8.1, Menlo Park, CA, USA, 2015) that measures biolayer interferometry (BLI). All steps were performed at 30 °C with shaking at 1500 rpm in a black 96-well plate containing 200 µL solution in each well. Streptavidin-coated biosensors were loaded with biotinylated peptides Bio-P1 or Bio-P2 (5 µg/mL) for 300 s followed by a wash. The sensors were then reacted for 300 s with increasing concentrations of antibody and then moved to buffer-containing wells for another 300 s (dissociation phase). Binding and dissociation were measured as changes over time in light interference after subtraction of parallel measurements from unloaded biosensors. Sensorgrams were fitted with a 1:1 binding model using the Octet data analysis software 8.1 (Fortebio, Menlo Park, CA, USA, 2015), and the presented values are an average of several repeated measurements.

### Western-blot

Chimeric MAb and rabbit sera were tested for specificity by western blot. Accordingly, SiMa cells (5×10^4^ cells/well) were cultured in serum-free Minimum Essential Medium (MEM) with Earls salts and Glutamax supplemented with N-2 (x1), B-27 (x1), and 25 mg/ml GT1b, as previously described [24], and differentiated for 48 hours in 96-well plates coated with poly-D-lysine. Cells were then exposed to 0, 400 or 4000 LD_50_/ml of BoNT/E, washed after 24 hours, and lysed by incubation with a cold solution of 0.1% Triton X-100, 150 mM NaCl, 1.5 mM MgCl2, 1 mM EGTA, 50 mM HEPES, and protease inhibitor cocktail (EDTA-free cOmplete tabs, Roche Diagnostics GmbH, Manheim, Germany) in water for 30 minutes. Lysates were loaded to a NuPAGE 10% Bis-Tris Gel (Invitrogen, CA, USA) and electrophoresed proteins were transferred to an Amersham Protran Premium 0.45 µm nitrocellulose membrane (GE Healthcare 10600096). The membrane was then blocked (1% skim milk in PBS) and probed with the chimeric MAb or immunized rabbit sera. Rabbit anti-SNAP-25 polyclonal antibody raised against a synthetic peptide corresponding to the N-terminus of human SNAP-25 (Sigma S9684), was used as a positive control to detect both cleaved and intact SNAP-25. AP-conjugated anti-human IgG (Sigma A3187) and AP-conjugated anti-rabbit IgG (Jackson 711-066-152) were used as secondary antibodies for chimeric MAb and rabbit sera, respectively. 5-Bromo-4-Chloro-3-Indolyl-phosphate (BCIP, Sigma B8503) was used as the chromogenic substrate.

## Results

### Immunization and characterization of the elicited antibody response

Successful isolation of specific high affinity antibodies from phage-display “immune-libraries” depends largely on the effectiveness of the immunization regimen. To elicit a polyclonal antibody response that will enable specific recognition of BoNT/E cleaved SNAP-25, mice and rabbits were immunized with an eight-residue synthetic peptide, coupled to KLH via an added cysteine residue (Table 1 and Fig1A), representing the resulting C-terminal end of SNAP-25 after BoNT/E activity. The immunization process included a prime injection and two monthly boosts, followed by two additional injections, given after two and three months respectively (Fig. 1B). Antibody titers were determined by ELISA against the specific synthetic peptide (Bio-P1), representing the resulting cleavage site of SNANP-25 after BoNT/E activity. Interestingly, while all immunized animals developed a significant antibody titer against the Bio-P1 peptide, the rabbits’ response was about 10 fold higher than that of the mice (Fig. 1C). Specificity was determined by comparing the response on the specific peptide (Bio-P1) to that attained on the non-specific peptide (Bio-P2), representing the non-cleaved SNAP-25 (Fig. 1C). Out of the five immunized mice, only one (m4) developed a specific response. In marked contrast, the two rabbits developed a highly specific antibody response toward Bio-P1, with no cross reactivity observed on Bio-P2. As the two rabbits’ antibody titers were similar to each other and about 10-fold higher than mouse m4, we decided to continue characterizing the rabbits’ sera.

**Figure 1:**
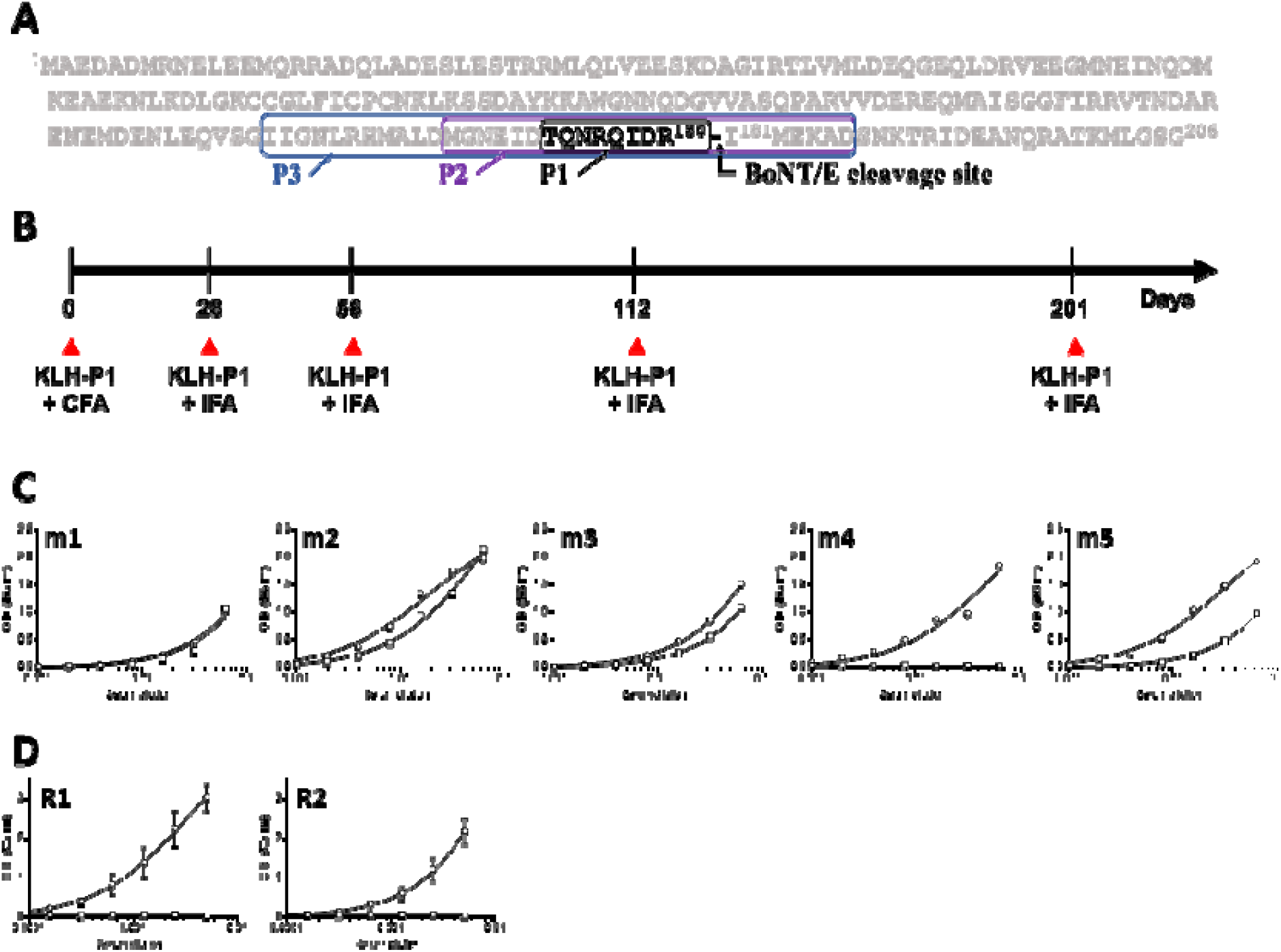
Peptide design, rabbit immunization and antibody serum levels. **A**. SNAP-25 protein sequence highlighting BoNT/E cleavage site and the peptides used in this study. **B**. Animal immunization regimen using KLH-P1 in either CFA (priming) or IFA (boosting). ELISA binding curves of serum of (**C**) immunized mice or (**D**) rabbits against Bio-P1 (circles) or Bio-P2 (squares). Lines represents non-linear regression fit and in (**D**) points are the mean ± STD of duplicates.

While specific recognition of the “cleaved” peptide was demonstrated, it was important to further verify that these antibodies can also specifically recognize the BoNT/E mediated cleaved SNAP-25. We previously demonstrated that SNAP-25 cleavage by BoNT/A can be monitored in intoxicated SiMa cells, using a specific antibody [25]. Here, cells were first incubated with 1000 LD_50_/ml of BoNT/E, washed and lysed 24 hours later. The cell lysate was resolved on SDS-PAGE and probed with an antibody directed against the N-terminus of the human SNAP-25. As expected, this antibody recognized both the intact (25 kDa) and the cleaved (23 kDa) SNAP-25 (Fig. 2). Next, the serum of each immunized rabbit was reacted with lysates of cells that were exposed to either 400 or 4000 LD_50_/ml of BoNT/E. Indeed, the sera of both animals specifically recognized only the cleaved SNAP-25, even at 400 LD_50_/ml, whereas no interaction was observed in the control cells. Taken together, these results clearly indicate that the immunization process elicited a strong and specific response, thus making the rabbits promising candidates for the isolation of the desired monoclonal antibodies.

**Figure 2:**
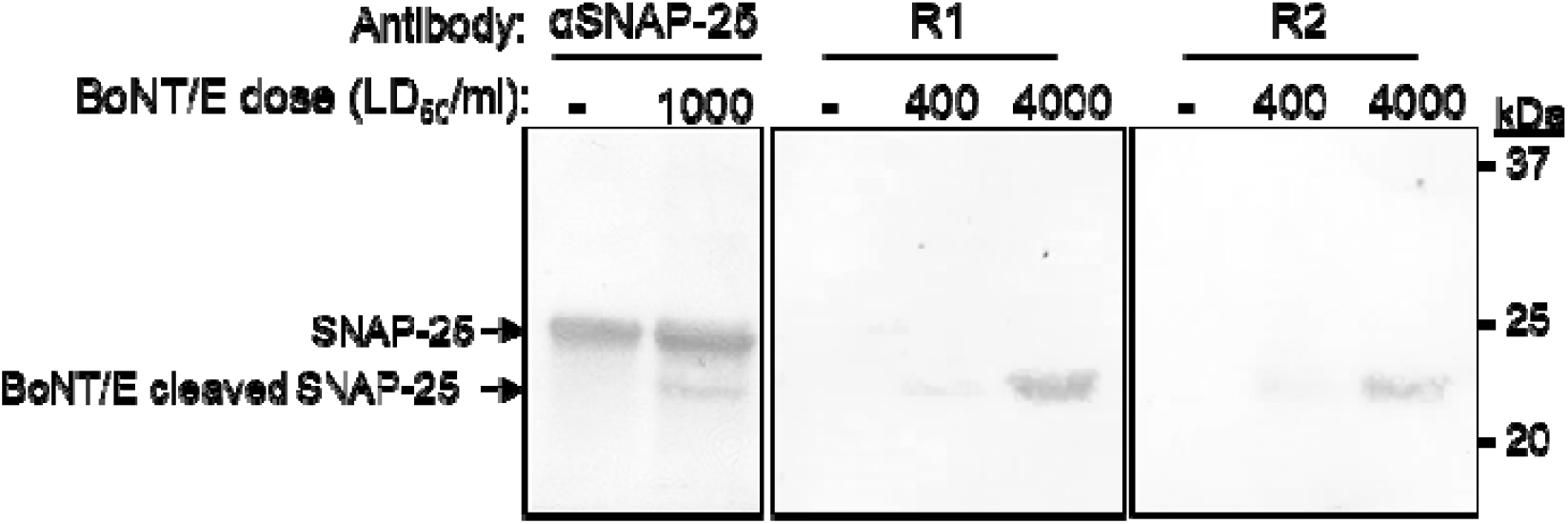
Specificity of the immunized rabbit’s sera. Differentiated SiMa cells were exposed to 0, 400 or 4000 LD_50_/ml of BoNT/E and lysed 24 hours after intoxication. Lysates were subjected to SDS-PAGE and Western-blotted using either anti-SNAP-25 polyclonal antibody or Bio-P1 immunized rabbits (R1 and R2).

### Library Construction and Panning

To ensure the highest level of antigen specific B cells, rabbits were sacrificed 10 days after the last boost [34-37]. A set of degenerate primers [29] was used to amplify rabbit VH and V_κ_ sequences from the spleen, bone marrow and peripheral blood of both rabbits. The VH and V_κ_ gene pools were then assembled by PCR to obtain combinatorial scFv fragments, which were than inserted into a phagemid vector resulting in the construction of two large (3×10^8^ pfu) phage-display libraries. To maximize the possibility of isolating a variety of antibodies, the libraries were subjected (separately) to three enrichment (panning) rounds, using streptavidin plates coated with Bio-P1 (Table 1). An additional enrichment course, carried-out with the combined libraries, included a depletion step, where a synthetic peptide (P3, Table 1), representing the intact SNAP-25, was diluted into the phage blocking buffer. At the end of the panning process, individual clones from each panning strategy were screened against adhered specific and non-specific peptides (Bio-P1 and Bio-P2 respectively, Table 1). Around 50% of the analyzed clones specifically recognized Bio-P1, for both enrichment processes involving the separate libraries. The 3^rd^ route, where a negative selection was applied, resulted in only 3% positive clones. Despite the different efficiencies, sequence analysis of the VH-V_κ_ genes revealed the emergence of only one antibody clone in each panning process (SNAP1, SNAP2, SNAP3). These were highly similar in their complementarity-determining regions (Table 2). Overall, the three antibodies shared 78% identical residues with additional 14% similar residues (conservative and semi-conservative homology) in their heavy chains and 87% identity and 8% similarity in their light chains. As the frameworks and CDRs of antibodies SNAP2 and SNAP3 shared the highest similarity, it is logical to assume that both originated from rabbit 1.

**Table 2:**
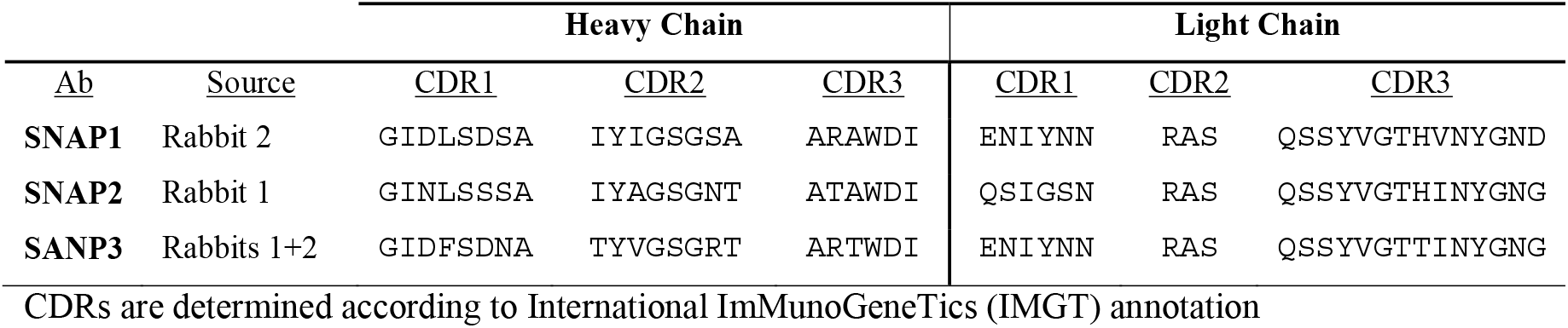
Amino-acid sequences of HCDRs and LCDRs.

Another interesting characteristic of all three isolated antibodies, is an uncommonly short heavy chain CDR3 (HCDR3, Table 3). While rabbit CDR3 heavy chains usually contain 12-13 residues [38, 39], the BoNT/E-cleaved SNAP-25 antibodies contain 6-residue CDR3s. This feature is nearly unique, as antibodies containing such short HCDR3 are expressed in less than 2.5% of sequenced rabbit antibodies [38, 39]. In addition, the apical region of the HCDR3 in rabbits is usually rich in glycine, erine and tyrosine, which are completely absent from this antibody. Instead, the HCDR3 contains a tryptophan, a bulky aromatic amino acid which is present in less than 2% of rabbit antibody HCDR3s [39]. To gain structural insight into this novel antibody, we used the AbPredict variable-domain structure prediction algorithm using default parameters [40] to generate three model structures (Fig. 3A). The three conformations are quite similar to one another and HCDR3 forms a groove, that probably mediates the specific binding to the cleaved SNAP-25. The model suggests that the tryptophan amino acid protrudes into the binding pocket where it may interact with the hydrophobic region of the neo-epitope (Fig. 3B).

**Figure 5:**
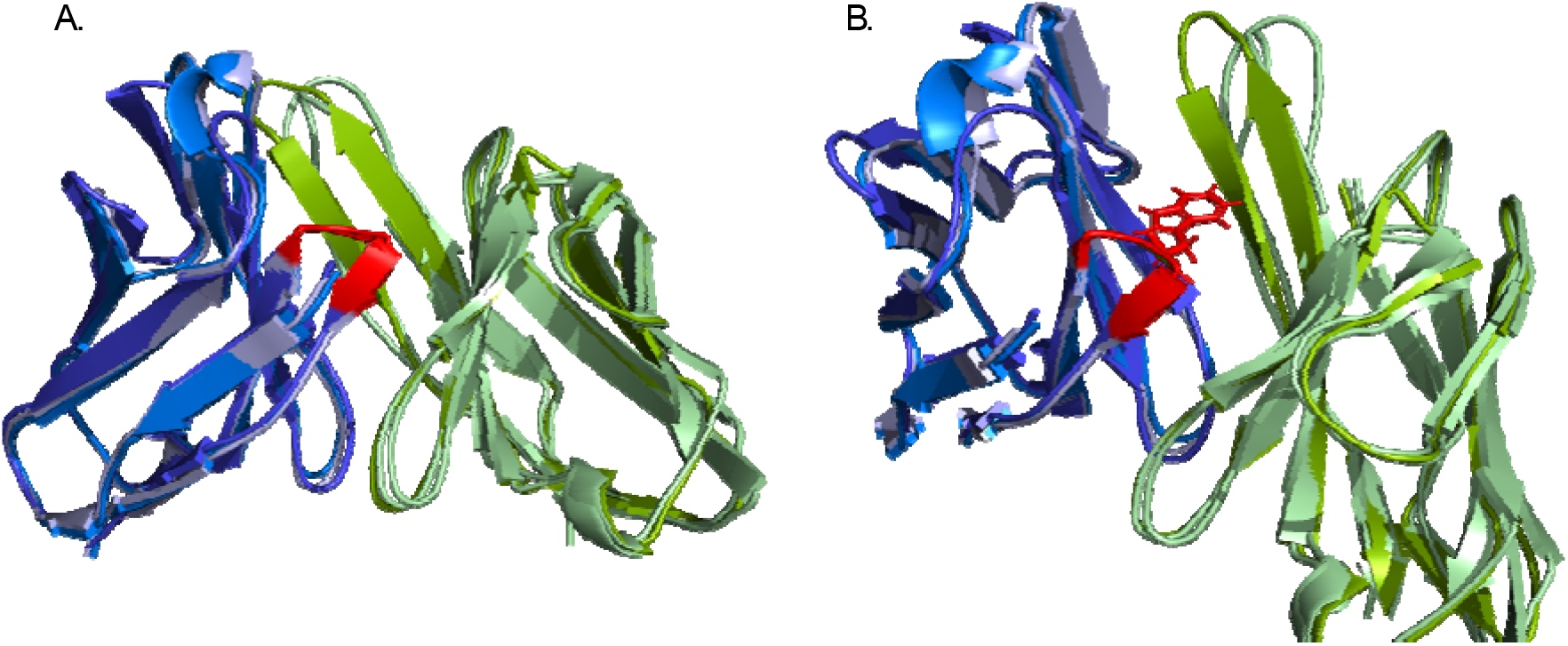
Superposition of the predicted structures of the variable regions of SNAP1, SNAP2 and SNAP3. Structure models were generated using AbPredict. Heavy chains are in blue shades, light chains are in green shades and HCDR3 is indicated in red. A. Ribbon representation of the three aligned antibody structures. B. The tryptophan is highlighted in sticks.

### Characterization of the Chimeric SNAP1 Antibody

Further characterization of binding and specificity requires a full-length antibody. As the three antibodies exhibited similar activities (determined by ELISA; unpublished data), SNAP1 was reformatted as a Fc-scFv antibody [29]. The specificity of the SNAP1 antibody was first demonstrated using ELISA, with plates coated with increasing concentrations of either the Bio-P1 or the Bio-P2 peptides. As expected, the antibody bound only the “cleaved” Bio-P1 peptide without any interaction with the Bio-P2 peptide (Fig. 4A).

**Figure 4:**
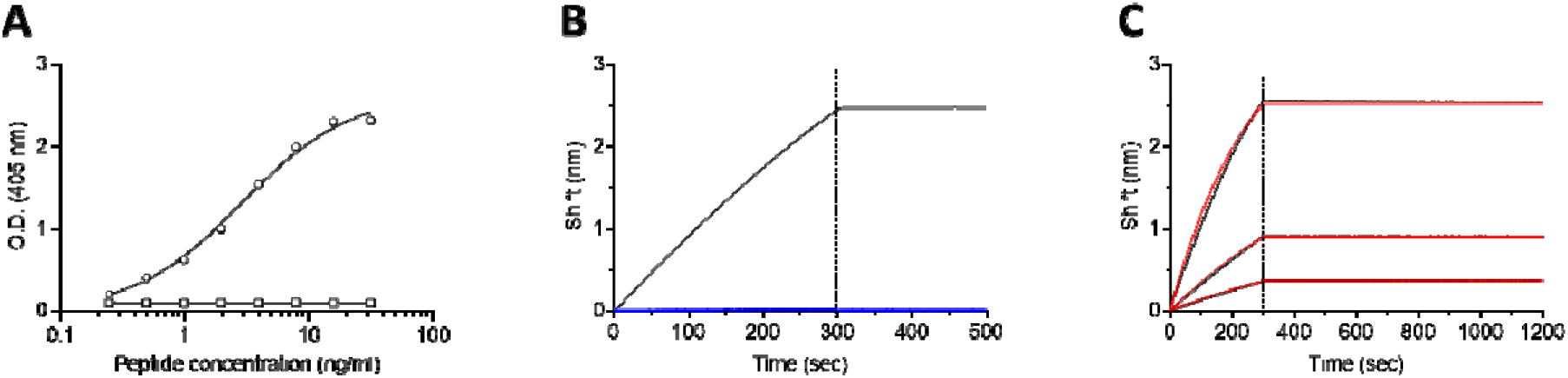
Specificity and affinity of SNAP1. **A**. Streptavidin-coated ELISA plates were loaded with increasing concentrations of either Bio-P1 (circles) or Bio-P2 (squares) peptides and the binding of SNAP1 antibody was determined. **B**. Octet Red BLI sensors were loaded with either Bio-P1 (black line) or Bio-P2 (Blue line) peptides. SNAP1 antibody was interacted with the sensors for 300 sec followed by wash (200 sec). **C**. Bio-P1 peptide was immobilized on a streptavidin-biosensor and reacted for 300 sec (association phase) with increasing concentrations of SNAP1 antibody (black lines; from bottom up: 6.7 nM, 20 nM, and 60 nM). The sensors were then immersed in buffer for another 900 sec (dissociation phase). Red lines: curve fitting of the 1:1 binding model.

The affinity of SNAP1 towards Bio-P1 was next determined using the Octet Red biolayer interferometry (BLI) system. Classical affinity measurements are generally carried out with a sensor-immobilized antibody (ligand) interacted with an analyte (peptide) in solution [31]. However, the use of small peptides as analytes is not recommended as they do not induce sufficient interference, thus limiting the accuracy of affinity measurments. We therefore immobilized the biotinylated peptides to the sensors and measured affinity using the antibody as the analyte. In order to minimize avidity effects, the peptides were loaded on the sensor ensuring low surface density (minimal wavelength shift, nm). To ensure assay specificity, the interaction of SNAP1 antibody was initially monitored against both Bio-P1 or Bio-P2 peptides. Indeed, the antibody specifically recognized the immobilized Bio-P1 and did not interact with Bio-P2 (Fig. 4B).

In order to determine SNAP1’s affinity towards cleaved SNAP-25, the Bio-P1 coated sensors were incubated with increasing concentrations of the antibody (the association phase) followed by a wash step (dissociation phase) (Fig. 4C). The sensorgrams were fitted with a 1:1 binding model, in order to determine the K_*on*_ and K_*off*_ rates. While the association rate of the antibody could be measured (5.8×10^5^ 1/Ms), the dissociation rate was extremely slow (below the Octet Red detection limit, 1×10^−7^ s^−1^) and could not be measured indicating that SNAP1’s affinity is in the sub-pM range.

### Selective detection of BoNT/E mediated cleavage of SNAP-25

As the major goal of this study was to develop a method that would enable the selective detection of the BoNT/E mediated cleaved SNAP-25 (SNAP-25_1-180_), we next tested whether the novel SNAP1 antibody is suitable for such an assay. Differentiated SiMa cells were exposed to BoNT/E at a concentration of either 400 or 4000 LD50/ml and SNAP-25 and the cell lysates were analyzed by SDS-PAGE. As a positive control, the anti-SNAP-25 antibody directed against the N-terminus protein region was applied and demonstrated a dose dependent cleavage of SNAP-25 by the toxin (Fig. 5). Next, cell lysates were incubated with SNAP1 antibody and indeed, a selective recognition of only the newly formed SNAP-25_1-180_ was achieved. To further demonstrate the specificity of the novel antibody, SiMa cells were exposed to BoNT/A which cleaves SNAP-25 between residues 197 and 198 (rather than between residues 180 and 181 by BoNT/E). The intoxication process by BoNT/A was verified using the anti-SNAP-25 N-terminus antibody, demonstrating the appearance of the expected cleaved product (Fig. 5). However, the BoNT/A cleaved SNAP25 could not be detected by the SNAP1 antibody, attesting to its high specificity toward the cleaved product of BoNT/E.

**Figure 5:**
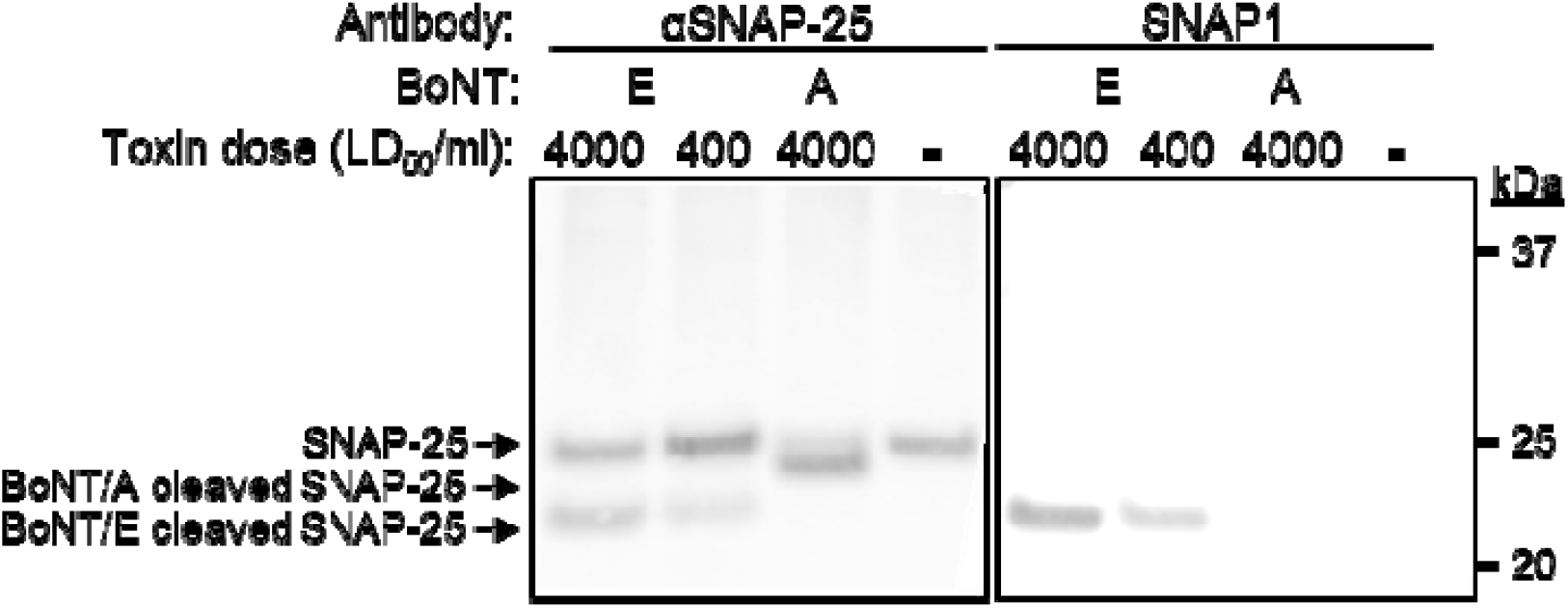
Specific detection of cleaved SNAP-25 neoepitope by SNAP1 antibody. Differentiated SiMa cells were exposed to the indicated doses of either BoNT/A or BoNT/E and lysed 24 hours after intoxication. Lysates were subjected to SDS-PAGE and Western-blotted using either anti-SNAP-25 polyclonal antibody or SNAP1.

## Discussion

In this work we describe the isolation of three scFv antibodies directed against a peptide that mimics the product of SNAP-25 following the enzymatic activity of BoNT/E. One of these clones (SNAP1) was further reformatted into a scFv-Fc antibody and was shown to bind the peptide with extremely high affinity and to specifically recognize the product of BoNT/E cleaved SNAP-25.

In the course of this study we immunized both mice and rabbits, aiming to broaden antibody repertoire. Results indicated that the humoral response of the immunized rabbits was both more robust and more specific than that observed in the immunized mice. Several reports have suggested that rabbits can develop antibodies against unique epitopes on human antigens that are not immunogenic in rodents [41] and to elicit strong immune responses against small molecules and haptens, which is rare in rodents [42-46]. These results may stem from the fact that rabbits belong to the taxonomic order Lagomorpha which is evolutionary distinct from the order Rodentia to which mice belong [47].

Another interesting finding was that the three isolated clones, originating from two different rabbits, share very high similarity in their sequence and structure. These unique features, including the short HCDR3 and the presence of the Trp amino acid in the HCDR3, most probably contribute to the high affinity and specificity of the SNAP1 antibody. Ig-blast analysis of these three clones revealed that their heavy chains share a common VH germline origin (IGHV1S69-1). In a previous work that performed next-generation sequencing (NGS) of the antibody repertoire in bone marrow and spleen of a naïve rabbit revealed that this germline is quite rare (0.01% or 0.05% respectively) [38]. However, in the same study, following immunization using a KLH-conjugated 16-mer peptide, the same germline gene was found to comprise 40% of the rabbit’s antibody repertoire [38]. In addition, a study of 235 cDNA clones from human peripheral blood suggested that a smaller CDR3 may create a unique “binding pocket” that allows better interactions between the antigens and CDRs 1 and 2 (Rosner Immunology 2001). It should also be noted that a similar, uncommonly short HCDR3 was observed in a mouse monoclonal antibody that specifically recognizes the cleavage site of SNAP-25 by BoNT/A [48]. As with the rabbit antibodies, this antibody contained a 5-residue HCDR3, whereas mouse antibodies most frequently contain 9-residue HCDR3s (6-residues HCDR3s manifest in less than 1% of the antibodies) [39]. Thus, it is suggested that the unique features of the SNAP1 antibody, namely its extremely high affinity and specificity towards the cleaved peptide, are derived from tight interactions in the formed unique pocket.

To summarize, by combining immunization and selection procedures we have isolated a novel, specific and highly potent antibody against the BoNT/E derived SNAP-25 neoepitope. This antibody can be further used to develop an in vitro cell-based assay to diagnose BoNT/E intoxication and to characterize antitoxin preparations, thus eliminating the use of animals in the standard mouse bioassay.

## Acknowledgments

We wish to express our gratitude to our colleague Dr. Itai Glinert for his fruitful remarks and helpful reviewing of this manuscript.

## References

1. Paddle, B.M., Therapy and prophylaxis of inhaled biological toxins. J Appl Toxicol, 2003. 23(3): p. 139–70.

2. Pirazzini, M., et al., Botulinum Neurotoxins: Biology, Pharmacology, and Toxicology. Pharmacological Reviews, 2017. 69(2): p. 200–235.

3. Lacy, D.B. and R.C. Stevens, Sequence homology and structural analysis of the clostridial neurotoxins. J Mol Biol, 1999. 291(5): p. 1091–104.

4. Simpson, L.L., Identification of the major steps in botulinum toxin action. Annu Rev Pharmacol Toxicol, 2004. 44 :p. 167-93.

5. Montal, M., Botulinum Neurotoxin: A Marvel of Protein Design. Annual Review of Biochemistry, 2010. 79(1): p. 591–617.

6. Rusnak, J.M. and L.A. Smith, Botulinum neurotoxin vaccines: Past history and recent developments. Hum Vaccin, 2009. 5(1 :(2p. 794-805.

7. Smith, L.A. and J.M. Rusnak, Botulinum Neurotoxin Vaccines: Past, Present, and Future. 2007. 27(4): p. 303–318.

8. Schiavo, G., M. Matteoli, and C. Montecucco, Neurotoxins affecting neuroexocytosis. Physiological Reviews, 2000. 80(2): p.717-766.

9. Binz, T., et al., Proteolysis of SNAP-25 by types E and A botulinal neurotoxins. J Biol Chem, 1994. 269(3): p. 1617–20.

10. Schiavo, G., et al., Botulinum neurotoxins serotypes A and E cleave SNAP-25 at distinct COOH-terminal peptide bonds.FEBS Lett, 1993. 335(1): p. 99–103.

11. Wang, J., et al., Novel chimeras of botulinum neurotoxins A and E unveil contributions from the binding, translocation, and protease domains to their functional characteristics. J Biol Chem, 2008. 283(25): p. 16993.7002-

12. Dembek, Z.F., L.A. Smith, and J.M. Rusnak, Botulism: cause, effects, diagnosis, clinical and laboratory identification, and treatment modalities. Disaster Med Public Health Prep, 2007. 1(2): p. 122–34.

13. MMWR, Investigational heptavalent botulinum antitoxin (HBAT) to replace licensed botulinum antitoxin AB and investigational botulinum antitoxin E. Morb Mortal Wkly Rep, 2010. 59(10): p. 299.

14. Division of Communicable Disease Control, C.D.o.P.H. Infant botulism treatment and prevention program. Available from: http://www.infantbotulism.org.

15. European Directorate for the Quality of Medicines and Healthcare, Botulinum toxin type A / type B for injection. 10 ed. European Directorate for the Quality of Medicines and Healthcare. Vol. EDQM Council of Europe. 2019, Strasbourg, France: European Pharmacopoeia.

16. Torgeman, A., et al., Studying the differential efficacy of postsymptom antitoxin treatment in type A versus type B botulism using a rabbit spirometry model. Dis Model Mech, 2018. 11.(9)

17. Behrensdorf-Nicol, H.A., et al., In vitro potency determination of botulinum neurotoxin serotype A based on its receptor-binding and proteolytic characteristics. Toxicol In Vitro, 2018. 53: p. 80–88.

18. Dressler, D. and G. Dirnberger, Botulinum toxin antibody testing: comparison between the immunoprecipitation assay and the mouse diaphragm assay. Eur Neurol, 2001. 45(4): p. 257–60.

19. Hanna, P.A. and J. Jankovic, Mouse bioassay versus Western blot assay for botulinum toxin antibodies: correlation with clinical response. Neurology, 1998. 50(6): p. 1624–9.

20. Lindsey, C.Y., et al., Evaluation of a botulinum fragment C-based ELISA for measuring the humoral immune response in primates. Biologicals, 2003. 31(1): p. 17–24.

21. Palace, J., et al., A radioimmuno-precipitation assay for antibodies to botulinum A. Neurology, 1998. 50(5): p. 1463–6.

22. Rosen, O., et al., Development of an Innovative in Vitro Potency Assay for Anti-Botulinum Antitoxins. Toxins (Basel), 2016. 8.(10)

23. Wild, E., et al., In vitro potency determination of botulinum neurotoxin B based on its receptor-binding and proteolytic characteristics. Toxicol In Vitro, 2016. 34: p. 97–104.

24. Fernandez-Salas, E., et al., Botulinum neurotoxin serotype A specific cell-based potency assay to replace the mouse bioassay. PLoS One, 2012. 7(11): p. e49516.

25. Torgeman, A., et al., An in vitro cell-based potency assay for pharmaceutical type A botulinum antitoxins. Vaccine, 2017.

26. Bak, N., et al., SiMa Cells for a Serotype Specific and Sensitive Cell-Based Neutralization Test for Botulinum Toxin A and E. Toxins (Basel), 2017. 9(7): p. 230.

27. Leveque, C., G. Ferracci, and e. al., An optical biosensor assay for rapid dual detection of Botulinum neurotoxins A and E. Scientific Reports, 2015 :5.p. 17953 DOI:10.1038/srep17953.

28. von Berg, L., et al., Functional detection of botulinum neurotoxin serotypes A to F by monoclonal neoepitope-specific antibodies and suspension array technology. Sci Rep, 2019. 9(1): p. 5531.

29. Mechaly, A., et al,.Novel Phage Display-Derived Anti-Abrin Antibodies Confer Post-Exposure Protection against Abrin Intoxication. Toxins (Basel), 2018. 10.(2)

30. Mechaly, A., et al., Inhibition of Francisella tularensis phagocytosis using a novel anti-LPS scFv antibody fragment. Scientific Reports, 2019. 9(1): p. 11418.

31. Noy-Porat, T., et al., Isolation of Anti-Ricin Protective Antibodies Exhibiting High Affinity from Immunized Non-Human Primates. Toxins (Basel), 2016. 8.(3)

32. Rosenfeld, R., et al., Isolation and Chimerization of a Highly Neutralizing Antibody conferring Passive Protection against Lethal B. anthracis Infection. PLoS ONE, 2009. 4(7)(7): p. e6351.

33. Bujak, E. and M. Matasci, eds. Reformatting of scFv antibodies into the scFv-Fc format and their downstream purification. Monoclonal antibodies: methods and protocols, ed. V. Ossipow and N. Fischer. Vol. 1131. 2014, Springer, New York.

34. Wrammert, J., et al., Rapid cloning of high-affinity human monoclonal antibodies against influenza virus. Nature, 2008 :453.p. 667–668.

35. Greenfield, E.A., Standard Immunization of Rabbits. Cold Spring Harb Protoc, 2020. 2020(9): p. 100305.

36. Overduin, L.A., J.J.M. van Dongen, and L.G. Visser, The Cellular Immune Response to Rabies Vaccination: A Systematic Review.Vaccines (Basel), 2019. 7.(3)

37. Webster, R.G., The immune response to influenza virus. II. Effect of the route and schedule of vaccination on the quantity and avidity of antibodies. Immunology, 1968. 14(1): p. 29–37.

38. Kodangattil, S., et al., The functional repertoire of rabbit antibodies and antibody discovery via next-generation sequencing. MAbs, 2014. 6(3): p. 628–36.

39. Lavinder, J.J., et al., Systematic characterization and comparative analysis of the rabbit immunoglobulin repertoire. PLoS One:(6)9.2014,p. e101322.

40. Norn, C.H., G. Lapidoth, and S.J. Fleishman, High-accuracy modeling of antibody structures by a search for minimum-energy recombination of backbone fragments. Proteins, 2017. 85(1): p. 30–38.

41. Rief, N., et al., Production and characterization of a rabbit monoclonal antibody against human CDC25C phosphatase. Hybridoma, 1998. 17(4): p. 389–94.

42. Feng, L., X. Wang, and H. Jin, Rabbit monoclonal antibody: potential application in cancer therapy. Am J Transl Res, 2011. 3(3): p.269-74.

43. Li, Y., et al., High affinity ScFvs from a single rabbit immunized with multiple haptens. Biochem Biophys Res Commun, 2000. 268(2): p. 398–404.

44. Liu, N., et al., Development of a new rabbit monoclonal antibody and its based competitive indirect enzyme-linked immunosorbent assay for rapid detection of sulfonamides. J Sci Food Agric, 2013. 93(3): p. 667–73.

45. Liu, N., et al., Simultaneous Raising of Rabbit Monoclonal Antibodies to Fluoroquinolones with Diverse Recognition Functionalities via Single Mixture Immunization. Anal Chem, 2016. 88(2): p. 1246–52.

46. Zhu, X., et al., Single-chain variable fragment (scFv) antibodies optimized for environmental analysis of uranium. Anal Chem, 2011. 83(10): p. 3717–24.

47. Miller, W., et al., 28-way vertebrate alignment and conservation track in the UCSC Genome Browser. Genome Res, 2007. 17(12): p. 1797–808.

48. Fernandes-Salas, E., J. Wang, and R. Kei, Immuno-based botulinum toxin serotype A activity assays. U.S. Patent no. US 12/403,531. (n. d.) Washington, D.C: U.S. Patent and Trademark Office, 2012.

